# The Drosophila small conductance potassium channel (SK) negatively regulates nociception

**DOI:** 10.1101/208868

**Authors:** Kia Walcott, Stephanie Mauthner, Asako Tsubouchi, Jessica Robertson, W. Daniel Tracey

**Author notes:** Corresponding author: W. Daniel Tracey Address: 702 N Walnut Grove Avenue, Bloomington, IN 47405. A present address: The University of Tokyo, Institute of Molecular and Cellular Biosciences, Neural Circuit Lab 1-1-1 Yayoi, Bunkyo-ku, Tokyo, 113-0032 JAPAN (Asako Tsubouchi).

## Abstract

Inhibition of nociceptor activity is important for the prevention of spontaneous pain and hyperalgesia. To identify the critical K^+^ channels that regulate nociceptor excitability we performed a forward genetic screen using a *Drosophila* larval nociception paradigm. Knockdown of three K^+^ channel loci, the *small conductance calcium-activated potassium channel* (*SK*), *seizure* and *tiwaz*, resulted in marked hypersensitive nociception behaviors. In more detailed studies of *SK*, we found that hypersensitive phenotypes could be recapitulated with a genetically null allele. Importantly, the null mutant phenotype could be rescued with tissue specific expression of an *SK* cDNA in nociceptors. Optical recordings from nociceptive neurons showed a significant increase in mechanically activated Ca^2+^ signals in *SK* mutant nociceptors. SK showed expression in peripheral neurons. Interestingly SK proteins localized to axons of these neurons but were not detected in dendrites. Our findings suggest a major role for SK channels in the regulation of nociceptor excitation and they are inconsistent with the hypothesis that the important site of action is within dendrites.

**Highlights:**

–Specific potassium channels regulate nociceptor excitability.
–SK channels have a critical function in nociception.
–SK channels specifically localize to sensory axons
–SK channels are not detectable in sensory dendrites.

## Introduction

The sensation of pain is important for avoiding exposure to noxious environmental stimuli that have the potential to cause tissue damage. These stimuli are detected by nociceptors, primary sensory neurons that detect noxious mechanical, noxious chemical and/or noxious temperatures. Transduction of noxious thermal, mechanical, and chemical stimuli is initiated by sensory receptor ion channels, which depolarize the sensory neuron plasma membrane and trigger action potentials (Dubin and Patapoutian, 2010). In the absence of such stimuli, healthy nociceptors remain relatively silent, with little spontaneous activity (Ritter and Mendell, 1992; Xiang et al., 2010) due to the action of potassium (K^+^) channels (Baumann et al., 2004) and chloride (Cl^−^) channels, which oppose depolarizing sodium (Na^+^) and calcium (Ca^2+^) currents. Despite their importance in keeping nociceptive neurons silent, the identity of the K^+^ channels that play the most critical roles in negatively regulating nociceptor excitability remains largely undetermined.

To identify these critical channels, we have carried out a forward genetic screen using a modified *Drosophila* larval nociception paradigm (Tracey et al., 2003; Zhong et al., 2010; Hwang et al., 2007; Zhong et al., 2012), which was optimized for the detection of hypersensitive nociception phenotypes. Using *in vivo* sensory neuron-specific RNA interference (RNAi) (Dietzl et al., 2007), we knocked down 35 of the predicted K^+^ channels that are encoded in the *Drosophila* genome (Table S1).

## Results and Discussion

### A paradigm to identify negative regulators of nociception

Studies performed in *Drosophila* have identified ion channel mutants with loss-of-function nociception insensitive phenotypes (*painless* (Tracey et al., 2003), *dTRPA1* (Walker et al., 2000; Kwan et al., 2006; Kernan et al., 1994; Zhong et al., 2012; Neely et al., 2011), *pickpocket* (Zhong et al., 2010; Hwang et al., 2007), *piezo* (Coste et al., 2012; Kim et al., 2012), *straightjacket* (Neely et al., 2010; Neely et al., 2011), and *balboa/ppk26* (Mauthner et al., 2014; Gorczyca et al., 2014; Guo et al., 2014). However, systematic screens to identify ion channels that specifically inhibit nociception have yet to be performed. The identification of loss-of-function nociception mutants that lead to hypersensitivity is desirable because these genes, by definition, function as negative regulators of nociception. Pharmacological agonists for negative regulators of nociception could represent potential therapeutics for the treatment of pain.

In previous studies that identified insensitive nociception mutants, larvae were contacted with a noxious heat probe of 46°C (Tracey et al., 2003; Hwang et al., 2007; Zhong et al., 2012; Zhong et al., 2010). This stimulus triggers a rapidly executed (within 1-2 seconds of contact) nocifensive escape locomotion (NEL) in wild type larvae. When performing NEL, the animal rotates (in a corkscrew-like fashion) around the anterior-posterior body axis. Noxious heat-insensitive mutants have been identified because they either failed to execute the NEL response, or because they showed a prolonged latency in their response, upon contact with the hot probe.

In contrast, near the threshold temperature for eliciting NEL (42°C) larvae require a relatively prolonged stimulus (greater than 6 seconds) to elicit a response (Figure 1). Therefore stimulation with a 42°C probe would identify hypersensitive nociception mutants that display a more rapid behavioral response to this threshold stimulus. Indeed this assay has been successfully used to identify genes required for hypersensitive thermal nociception (Honjo et al., 2016).

**Figure 1.**
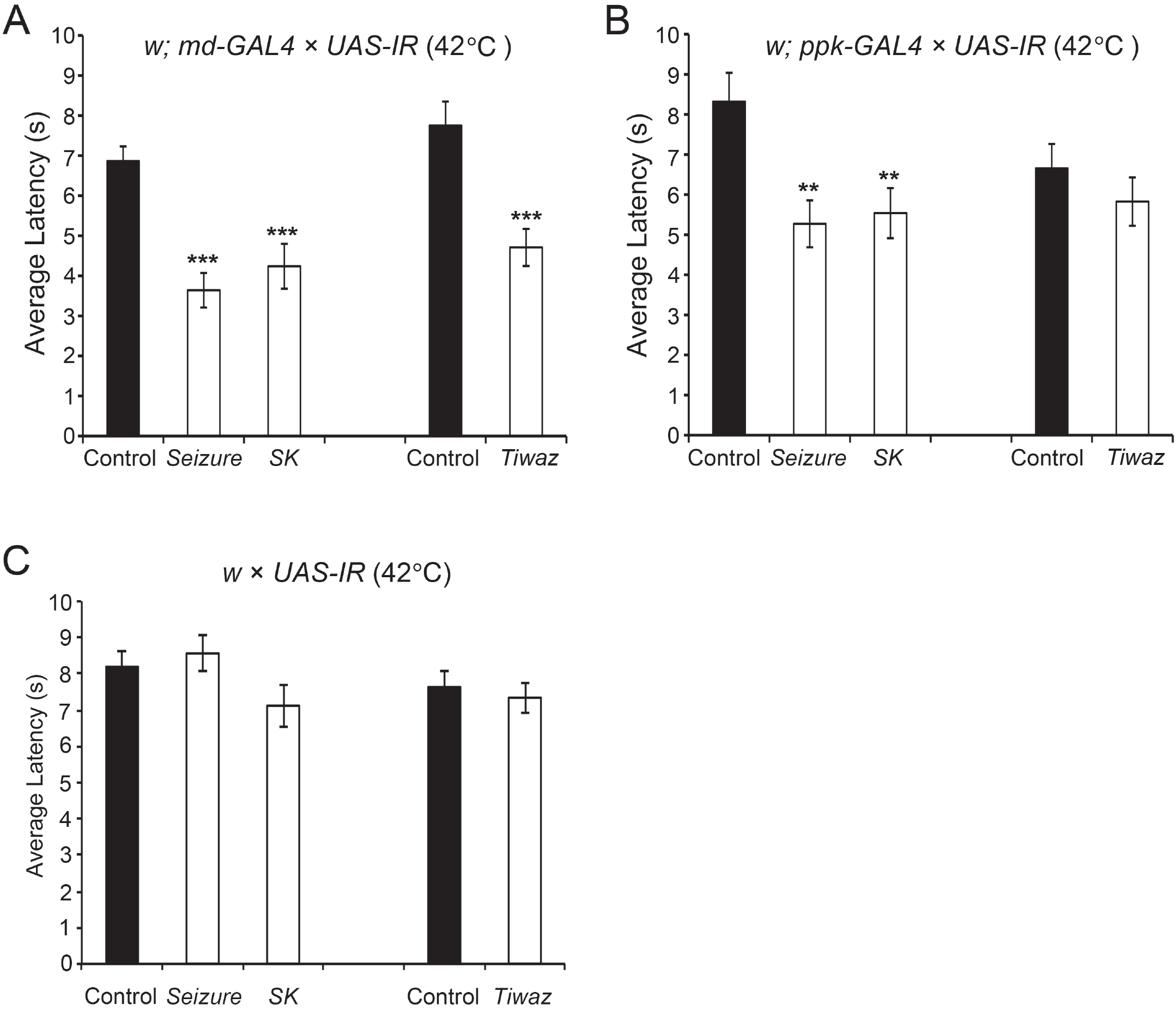
Thermal nociception responses of K^+^ channel knockdown larvae. (A) In response to a threshold noxious temperature (42°C), *UAS-seizure-IR* (VDRC strain P{KK105733}v104698)(n = 33) and *UAS-SK-IR* (VDRC strain P{KK107699}v103985) larvae (n = 25) show reduced latency when crossed to *md-Gal4; UAS-Dicer2* compared to control *yw; attP* strain (genotype: *yw/w; md-GAL UAS-Dicer2/attp{empty}*) (n = 70). One-way ANOVA with Dunnett’s Test, P< .001 = ***. *tiwaz-*RNAi (TRiP strain P{TRiP.JF01867}attP2) larvae (n = 40) show hypersensitivity to noxious heat compared to control *yv; attP* animals (genotype: *yv/w; md-GAL UAS-Dicer2/+; attp2{empty/+}*) (n = 37 Student’s T-test, P< .05 = *). (B) The average latency of the K^+^ channel UAS-RNAi lines targeting *SK*, *seizure*, and *tiwaz* crossed to *ppk-Gal4; UAS-Dicer2* (42°C). *SK* (n = 34) and *seizure* (n = 35) RNAi animals show reduced latency to perform nociception behaviors compared to control *yw; attP* animals (genotype: *yw/w; ppk-GAL UAS-Dicer2/attp{empty}*) (n = 31). One-way ANOVA with Dunnett’s Test, P< .05 = *. *tiwaz-*RNAi animals (n = 44) do not show significant hypersensitivity to noxious heat compared to control *yv; attP* animals (genotype: *yv/w; ppk-GAL UAS-Dicer2/+; attp2{empty/+}*) (n = 54) Student’s T-test, P = 0.29. Error bars indicated SEM. (C) RNAi transgenes alone do not show thermal nociception defects. RNAi transgenes targeting *SK*, *seizure*, and *tiwaz* were each crossed to *w*^1118^ strain. In the thermal nociception assay (42°C), no difference was detected between RNAi alleles targeting *SK* and *seizure* in the absence of a driver (n = 32 and n = 25) compared to the control strain (*yw;attP*) (n = 40) One-way ANOVA with Dunnett’s Test. Similarly, animals carrying the *tiwaz*-RNAi transgene in the absence of the driver (n = 41) are not different from the control strain (*yv;attP*) (n = 38) Student’s T-test, P = .63. Error bars indicate SEM.

We performed a forward genetic screen in which we tested the effects of RNA interference (RNAi) knockdown of the predicted potassium (K^+^) channels in the md neurons. Since the general function of K^+^ channels is to promote hyperpolarization of neuronal membranes, these channels are expected to inhibit neuronal firing.

### Tissue-specific knockdown of *Drosophila* genes *in vivo*

A collection of transgenic RNAi strains has been constructed by the Vienna *Drosophila* RNAi Center (VDRC) (Dietzl et al., 2007) and the Transgenic RNAi Project (TRiP). These strains allow for tissue-specific gene silencing *in vivo* under control of the Gal4/UAS system (Dietzl et al., 2007; Brand and Perrimon, 1993; Ni et al., 2009). We identified 54 UAS-RNAi lines in these collections that both targeted 35 K^+^ channels and which had few predicted off target effects (Table S1). All K^+^ are represented in our assembled collection.

We first investigated the effects of knocking down the K^+^ channels under control of the GAL4 109(2)80;UAS-*Dicer2* (*md-Gal4*;*UAS-Dicer2*) driver strain. This strain drives UAS transgene expression in the Class I, II, III and IV md neurons (Gao et al., 1999). Evidence suggests that the major nociceptive function is mediated by the class IV md neurons but Class II and Class III neurons are also involved (Hwang et al., 2007; Hu et al., 2017). The use of *UAS-Dicer2* in the driver strain results in more efficient gene silencing (Dietzl et al., 2007). To perform the screen the *md-Gal4*;*UAS-Dicer2* driver strain was crossed to each of the 54 UAS-RNAi strains targeting the K^+^ channels and the nocifensive responses of the larval progeny stimulated with a 42°C heat probe were observed. The crossed progeny from three UAS-RNAi lines targeting three distinct K^+^ channel subunits showed a significantly more rapid response relative to genetic background control strain. These UAS-RNAi lines targeted the *Small Conductance Calcium Activated Potassium Channel* (*SK)* (Abou Tayoun et al., 2011), the *seizure* channel (in the *ether-a-gogo related* family) (Jackson et al., 1984), and the *tiwaz* gene (Williams et al., 2014) (encodes a protein with homology to the potassium channel tetramerization domain) (Figure 1 A,C). Although phenotypes were not observed for other tested K^+^ channels, this method for RNAi is prone to false negatives so potential involvement for other channels cannot be ruled out by our screen. To test whether the effects of the RNAi were specific to the nociceptive class IV sensory neurons we next tested animals expressing UAS-RNAi targeting these three candidates under control of *ppk-Gal4*;*UAS-Dicer2* (Ainsley et al., 2003). The hypersensitive responses persisted in *SK-RNAi* and *seizure*-RNAi animals (Figure 1 B,C).

### SK negatively regulates thermal nociception behavior

Recent work on dissociated mammalian DRG primary sensory neurons showed that the SK inhibitor apamin increased the action potential frequency of mammalian nociceptors (Pagadala et al., 2013). While this work suggested an SK-mediated pathway in establishing DRG neuron excitability, the cellular role of this ion channel specifically in nociception remains largely unexplored and has not been verified with genetic mutants. The mammalian genome contains three genes that encode SK channel subunits (Köhler et al., 1996) while the *Drosophila* genome encodes only a single *SK* locus on the X chromosome at the cytological position 4F5-4F9 (Abou Tayoun et al., 2011; Adelman et al., 2012; Köhler et al., 1996). The locus is 60kb in length and is predicted to encode at least fourteen distinct transcripts (Gramates et al., 2017) (Figure 2A (for simplicity only a single transcript is shown)). The *Drosophila SK* locus has been found to mediate a slow Ca^2+^ activated K^+^ current in photoreceptor neurons and muscle as well as playing a role in learning and memory (Abou Tayoun et al., 2011; Abou Tayoun et al., 2012; Gertner et al., 2014). To further investigate the function of SK, we generated a DNA null mutant by deleting the gene with Flippase (FLP) and FLP Recombination Target (FRT) containing piggyBac transposons (Thibault et al., 2004; Parks et al., 2004) (Figure 2A). Deletion of the genomic region was confirmed by polymerase chain reaction (Figure S1A,B). Consistent with the hypothesis that SK is an important negative regulator of nociception, *SK* null mutants showed a pronounced hypersensitive response at 42°C. These animals showed an average response latency to a 42°C stimulus of 3.2 seconds, which was significantly faster from than the parental strain (Exelixis isogenic *w*) response of 6.2 seconds (Figure 2B).

**Figure 2.**
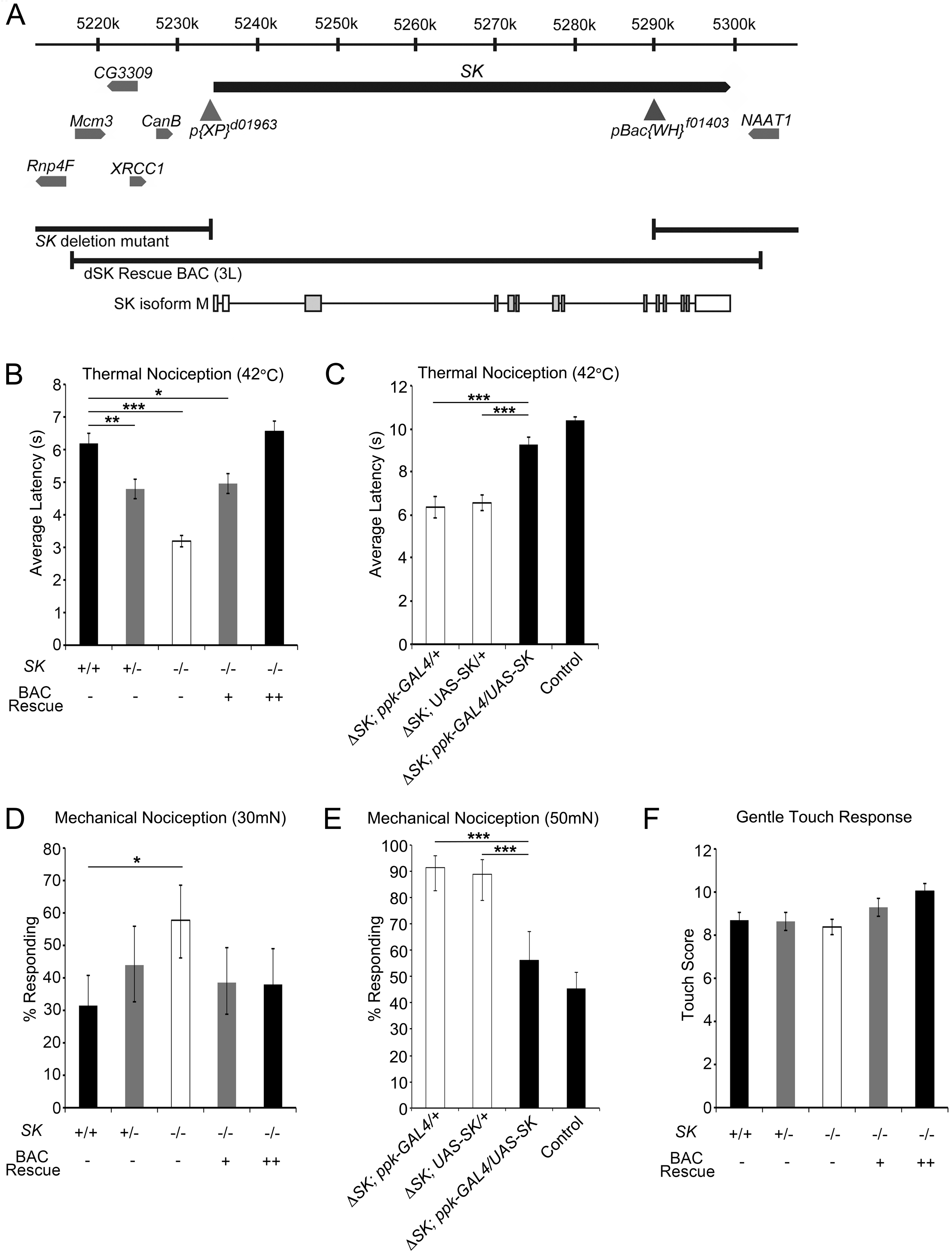
Expression of SK in class IV neurons could rescue the nociceptive phenotype. (A) Schematic representation of the genetic interval surrounding *SK* locus. The isoforms of *SK* and *SK* rescue BAC are schematically shown. The locations of *pBac* transposable elements used to generate *SK* null mutants are shown as gray triangles. Class IV specific expression of *SK-M* rescues the hypersensitive thermal phenotype in *SK* mutant larvae. (B) *SK* mutant animals are hypersensitive to noxious heat (42°C). *SK* null mutants (white bar, n = 66) show hypersensitivity to noxious heat compared to control animals (n = 66). Replacing *SK* in the genome with two copies of the BAC transgenes rescue the nociception defect (black bar, n = 66). The *SK* mutant hypersensitivity phenotype is semi-dominant as animals heterozygous for the deletion (gray bars, n = 72) show hypersensitivity compared to control animals One-way ANOVA with Dunnett’s Test, P< .001 = ***, P< .01 = **, P< .05 = *. Error bars indicate SEM. (C) Latency to respond to a 42°C probe in larvae expressing *SK-M* in the class IV neurons is significantly different from wild type control. Data are presented as mean ± SEM. *SK; ppk-GAL4/+* (*ppk-GAL4 only*, n = 45), *SK; UAS-SK(M)/+* (*UAS-SK only*, n = 72), *SK; ppk-GAL4/UAS-SK(M)* (Rescue, n = 49), and Exelixis isogenic white (Control, n = 70). One-way ANOVA followed by Tukey’s test. P< .001 = ***. (D) Mechanical nociception assay using a 30mN von Frey. *SK* null mutants (white bar, n = 71) show hypersensitivity to noxious mechanical stimulation compared to control animals (n = 81). Replacing *SK* in the genome with two copies of the BAC transgenes rescue the nociception defect (black bar, n = 79). The *SK* mutant hypersensitivity phenotype is semi-dominant as animals heterozygous for the deletion (gray bars, n = 66 and n = 83) show hypersensitivity compared to control animals. Fisher’s Exact Test with Holm-Bonferroni correction. Data are presented as percentages ± 95% confidence intervals. P< .05 = *. (E) Expression of SK isoform M in class IV neurons rescues the phenotype of *SK* mutant. *SK; ppk-GAL4/+* (*ppk-GAL4* only, n = 70), *SK; UAS-SK(M)/+* (*UAS-SK* only, n = 63), *SK; ppk-GAL4/UAS-SK(M)* (Rescue, n = 73), and Exelixis isogenic *white* (Control, n=244). Fisher’s Exact Test with Holm-Bonferroni correction. Data are presented as percentages ± 95% confidence intervals. P< .001 = ***. (F) *SK* null mutant animals show normal gentle touch responses One-way ANOVA with Dunnett’s Test, P> .05. Errors bars indicate SEM.

To confirm that loss of SK was responsible for the nociception defect, we performed a genetic rescue experiment through transgenic insertion of an ~ 80kb bacterial artificial chromosome (BAC) transgene that covered the entire *SK* locus (Venken et al., 2006). The BAC transgenic flies were crossed into the *SK* genetic mutant background to create rescue animals containing either one or two copies of the BAC transgene covering the *SK* genomic region (Figure 2A). The nociception hypersensitivity phenotype was fully reverted in rescue animals containing two copies of the BAC transgene (Figure 2B). Interestingly, female animals heterozygous for the *SK* mutation also exhibit hypersensitivity to noxious heat but to a lesser degree than homozygotes (4.8 seconds). Furthermore, one copy of the BAC rescue transgene provides only partial rescue of the hypersensitivity phenotype (5.0 seconds) (Figure 2B). These data combined support a dosage sensitive semi-dominant thermal nociception defect for *SK* mutants that requires two copies of the BAC transgene for full rescue.

To test for a nociceptor specific requirement for *SK*, we expressed the *SK-M* transcript under control of the *ppk-GAL4* driver in the *SK* mutant background (Figure 2A). This manipulation fully rescued the hypersensitive nociception phenotype of the *SK* mutant animals (Figure 2C) confirming the site of action for SK in the nociceptor neurons.

### SK negatively regulates mechanical nociception behavior

The elaborately branched class IV neurons function as polymodal nociceptors, playing a role in both thermal (≥ 39°C) and mechanical nociception (≥ 30mN). Channels expressed in class IV neurons such as such Painless and dTRPA1 are required for both thermal and mechanical nociception while Pickpocket, Balboa/PPK26, and Piezo have more specific roles in mechanical nociception (Tracey et al., 2003; Zhong et al., 2010; Zhong et al., 2012; Kim et al., 2012; Mauthner et al., 2014). Interestingly, *SK* mutant larvae showed enhanced nocifensive responses to a 30mN mechanical stimulus compared to parental strain animals (Figure 2D). With this stimulus, SK null mutant animals respond to 58% of the noxious force stimuli while control animals respond to 31%. As with thermal nociception, replacing *SK* in the genome by BAC transgene restored the mechanical nociception response to wild-type levels with 38% of BAC rescue animals responding to the 30mN stimulus (Figure 2D). However, the mechanical nociception phenotype was less sensitive to dosage. Animals heterozygous for the SK null mutation as well as animals containing one copy of the BAC rescue transgene respond similarly to wild-type animals (44% and 39%, respectively). As with thermal nociception, *UAS-SK-M* expressed under control of the *ppk-GAL4* driver fully rescued the *SK* mutant mechanical nociception phenotype (Figure 2E).

To determine if SK disruption affects mechanosensation in general, we tested *SK* mutant larvae in an established gentle touch assay (Tsubouchi et al., 2012)) (Kernan et al., 1994; Hwang et al., 2007; Zhong et al., 2010; Tracey et al., 2003). Gentle touch responses in *SK* mutants appeared normal (Figure 2F). Thus, the somatosensory effects of SK were more specific to the nociception pathway, regulating nociceptor activity both in response to noxious thermal and mechanical stimuli.

### SK negatively regulates nociceptor excitability

Next, we performed optical recordings upon control (Exelixis isogenic *white*) and *SK* null mutant larvae expressing the genetically encoded Ca^2+^ indicator, GCaMP3.0 (Tian et al., 2009), under the control of the nociceptor-specific driver, *ppk*-Gal4. In this filleted larval preparation, the md neurons expressing GCaMP3.0 were imaged through the transparent cuticle using high-speed, time-lapse confocal microscopy while being exposed to 50mN force stimuli. *ppk-Gal4* expressing neurons imaged in this preparation showed rapidly increasing GCaMP3.0 signals during the initial application of force and this signal rapidly declined (Figure 3A). In *SK* mutant animals, the peak calcium response (measured at the cell soma) was significantly increased relative to wild type and the signal remained elevated above the baseline for several seconds following the mechanical stimulus (Figure 3B). Restoration of SK-M to the mutant background rescued the elevated peak response but did not fully suppress the prolonged signal seen in the mutant (Figure 3B). However, the prolonged signal still appeared to be slightly elevated in the rescue animals (Figure 3B). Thus although SK-M can rescue behavioral phenotypes, and peak calcium responses in Class IV neurons, it is possible that one or more of the 13 other isoforms is required for complete restoration of wild type responses in this Ca^++^ imaging assay.

**Figure 3.**
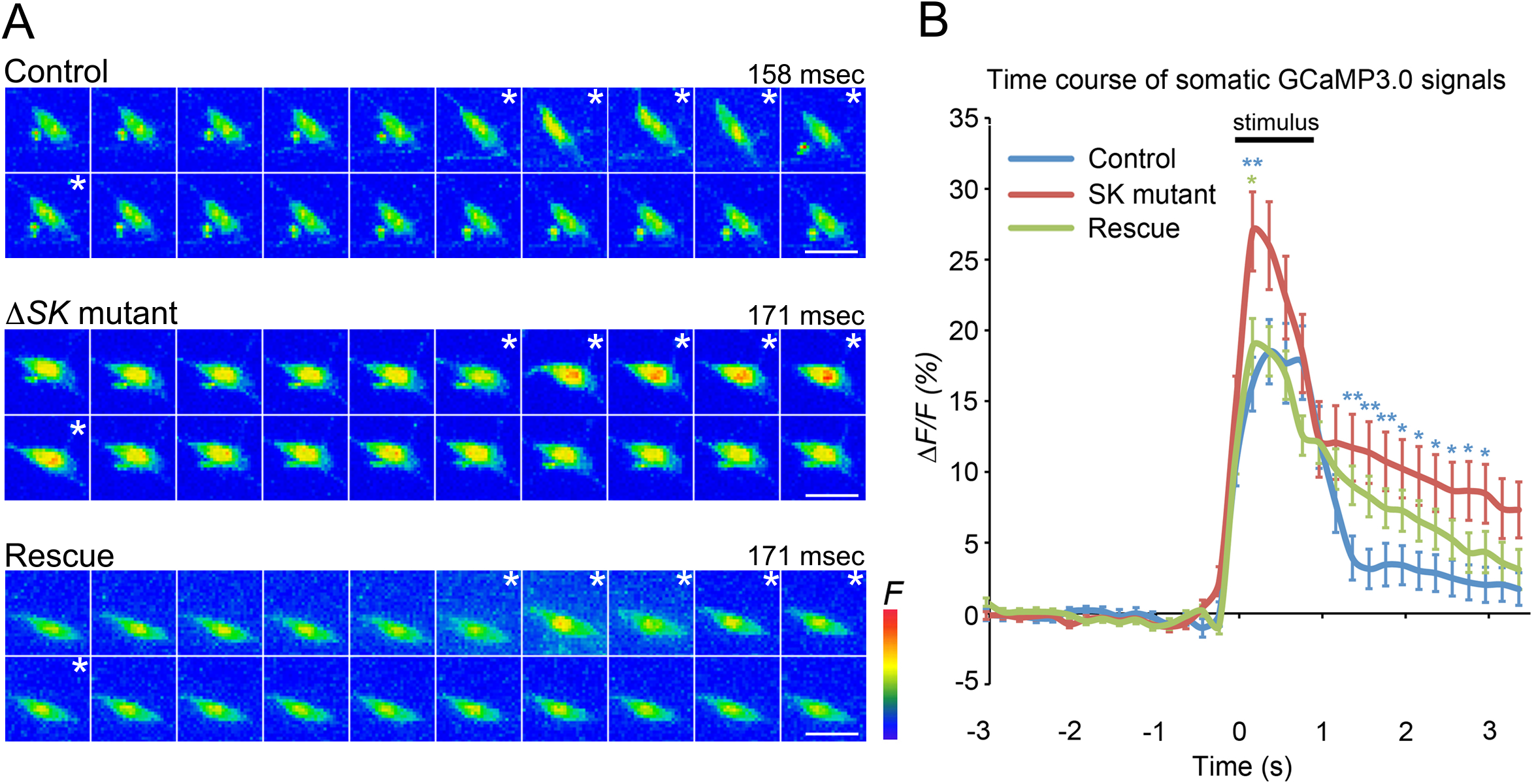
Optical recordings show *SK* mutant enhances force triggered Ca^2+^ response of class IV neurons. (A) Maximum intensity projections from Z-stack time series. (Control: upper) Images of representative class IV ddaC neuron expressing GCaMP3.0, displaying increased GCaMP3.0 fluorescence during stimulation (asterisks) (1 frame = 158 msec). (*SK* mutant: middle) Images of representative *SK* mutant ddaC neuron shows higher increase in GCaMP3.0 intensity relative to controls during stimulation (asterisks) (1 frame = 171 msec). (Rescue: bottom) Images of representative ddaC neuron repressing UAS-SK isoform M in *SK* mutant background. (1 frame = 171 msec). Scale bars = 10 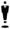 m. Genotypes were *w; ppk-GAL4 UAS-GCaMP3.0/+* (Control), *w SK^−^; ppk-GAL4 UAS-GCaMP3.0/+* (*SK* mutant), and *w SK^−^; ppk-GAL4 UAS-GCaMP3.0/UAS-SK(M)* (Rescue). (B) Traces showing average peak *∆F/F_0_* % of ddaC neurons in each genotype before, during, and after mechanical stimulation. The bar above the trace shows the average length of the mechanical stimulus (0.93 ± 0.02 sec). Statistical analysis was performed using a one-way ANOVA with Tukey’s HSD post-hoc for pair-wise comparisons. Data are means ± s.e.m. Blue asterisks show Control vs. *SK* mutant. Green asterisk shows *SK* mutant vs. Rescue. P< .01 = ** and P< .05 = * in each time point. Control (n = 50), *SK* mutant (n = 55), and Rescue (n = 59).

### The *SK* gene is expressed in class IV md sensory neurons

To evaluate the expression of *SK* we first employed a gene-trap strategy using a transgenic *Drosophila* strain from the *Minos-*mediated integration cassette (MiMIC) collection (Venken et al., 2011). A MiMIC element inserted in the proper orientation into the 5’ non-coding intron of SK long isoform transcripts expressed EGFP in the native pattern (i.e. at endogenous levels in appropriate tissues) of SK. We observed EGFP expression in the larval peripheral nervous system in a subset of type I and type II sensory neurons that included the class IV md neurons (Figure 4A). These results reveal that transcripts encoding long-isoform SK proteins are endogenously expressed in the nociceptors and provide additional validation of our tissue-specific *UAS-SK-M* rescue experiments (that restored normal nociception function to *SK* mutant larvae) (Figure 2 C,E).

**Figure 4.**
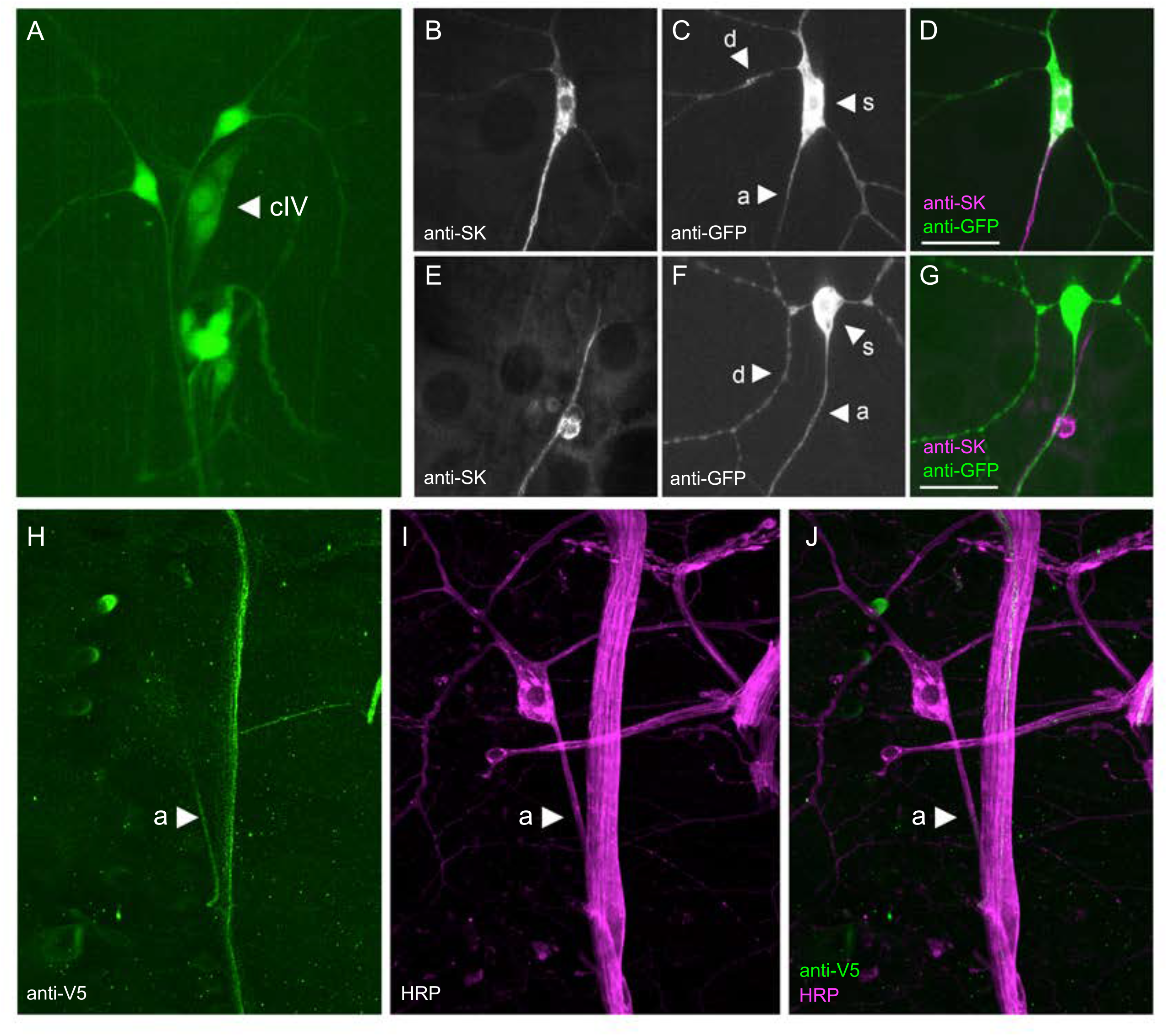
*SK* is expressed in class IV neurons and SK::V5 proteins localize at the axons. Maximum intensity projection from z-stack images representative of the dorsal md neuron cluster (segment A4-A6) in larvae. (A) A MiMIC element inserted in the 5’UTR of *SK* expresses GFP in multidendritic, bipolar, and external sensory neurons (genotype: *y^1^ w^*^ Mi(MIC) SK^MI10378^*). (B-D) Anti-SK antibodies label class IV expressed SK isoform M proteins in *SK* null mutant background (genotype: *SK^−^; ppk-GAL4 UAS-mCD8::GFP/UAS-SK-(M)*). SK protein expression (B and magenta in D) and class IV neuron (C and green in D). (E-G) Endogenous SK protein is detected in axons of sensory neurons in control larvae (genotype: *ppk-GAL4 UAS-mCD8::GFP/+*). SK protein expression (E and magenta in G) and class IV neuron (F and green in G). (H-J) Anti-V5 antibodies label class IV axons of *SK::V5* larvae (genotype: *w SK::V5*). d: dendrite, s: soma, and a: axon. Scale bars are 10μm.

### SK proteins are localized to a proximal compartment of sensory axons

The potassium currents mediated by SK channels in mammalian neurons make an important contribution to the after hyperpolarization (AHP) of action potentials (Faber and Sah, 2002; Pedarzani et al., 2005). However, the precise subcellular function of SK channels is thought to vary depending on the subcellular compartment where it resides. SK proteins have been found in somatic or dendritic compartments in some cell types (reviewed (Rudolph and Thanawala, 2015), and in axons or presynaptic compartments in others (Abou Tayoun et al., 2011). Thus, we wished to identify the subcellular compartment containing SK channels in the nociceptive md neurons.

Using a previously generated anti-SK antibody (Abou Tayoun et al., 2011) we first observed the localization of the SK-M protein that we used in our experiment for rescue of mutant phenotypes (Figure 4B-D). Surprisingly, we found that the rescuing SK-M protein was strongly detectable in the class IV axons and soma (Figure 4B,D) but only weakly detectable in proximal dendrites (Figure 4C,D). This was consistent with staining in wild type animals (Figure 4E-G), where we were able to detect SK protein in axons (Figure 4E,G) and soma of sensory neurons but we were unable to detect any expression in the dendrites or soma of class IV neurons (Figure 4F, G). The anti-SK staining was completely eliminated in null mutant animals confirming specificity of the antibody (Figure S1C-E).

These results raised the possibility that SK protein is required in axons and not in dendrites for nociception. However, the anti-SK antibody was raised against an N-terminal fragment of SK protein, and this domain of the protein is absent in 6 of the 14 predicted SK isoforms (Gramates et al., 2017). Thus, it remained possible that short isoforms of SK might localize to dendrites. Thus, in order to detect as many SK proteins as possible, and at their endogenous expression levels, we used CRISPR mediated homologous repair (Gratz et al., 2013; Jinek et al., 2012; Cong et al., 2013) to introduce a V5 epitope tag (Hampel et al., 2011) at the *SK* locus (Figure S2A-C). The inserted V5 epitope tag is encoded by an exon present in 13 of the 14 known *SK* transcripts (with the exception of the *SK-J*) located immediately downstream of the sequence encoding the SK calmodulin-binding domain (Figure S2A-A’). Interestingly, anti-V5 directed immunofluorescence in animals with our genomic modification was present in the axons of peripheral sensory neurons. Specifically, we observed strong labeling of axons of a subset of type I (external sensory and chordotonal; data not shown) neurons and type II multidendritic neurons, including the class IV (Figure 4H-J). Interestingly SK::V5 proteins concentrate within a proximal compartment of axons in class IV sensory neurons (Figure 4H,J) but were not detected in nociceptor axon terminals in the CNS (Figure S3A-C’) or in sensory dendrites of the nociceptive neuron arbor (not shown). The subcellular distribution of SK::V5 proteins along the axon of sensory neurons corroborated labeling of similar neuronal structures with anti-SK antibodies targeting long isoforms of SK (Abou Tayoun et al., 2011) (Figure 4E-G). Note that the SK-J protein contains the N-terminal epitope that is detected by the anti-SK making it unlikely that it localizes to dendrites. Thus, our evidence combined suggests SK proteins in axons, and not in dendrites are important for nociception. As with all antibody staining approaches, we cannot exclude the possibility that SK channels present in dendrites exist but are beneath the limits of detection by this approach.

In summary, *SK* mutants showed hypersensitivity in both thermal and mechanical nociception assays. The mammalian genome encodes three SK channels belonging to the KCNN gene family while the *Drosophila* genome encodes only one SK subunit (Abou Tayoun et al., 2011; Adelman et al., 2012; Köhler et al., 1996). In mammals, SK channels are widely expressed in the nervous system where they play an important role in the AHP of the action potential (Köhler et al., 1996; Stocker and Pedarzani, 2000; Tacconi et al., 2001; Hosseini et al., 2001). The SK channel blocker apamin is a component of bee venom that causes acute pain when injected into the skin of mammals and recent results support a role for SK channels in mediating an important inhibitory role in mammalian neurons (Ohtsuki et al., 2012; Pagadala et al., 2013).

Our finding on the axonal localization of SK channel proteins is interesting in several respects. First, it suggests that the enhanced Ca^++^ signals that we observed in nociceptor soma are potentially caused by back propagation of action potentials rather than a hyperexcitable soma or dendrites. Second, the proximal axonal localization is consistent with recently described evidence that other *Drosophila* K^+^ channels may be localized to the axon initial segment in these neurons (Jegla et al., 2016). It is possible that SK regulates the after hyperpolarization of action potentials as in other systems, and this in turn, regulates firing frequency. It has been proposed that bursts and pauses in firing of the *Drosophila* nociceptor neurons may be necessary for robust nociception responses (Terada et al., 2016). As well, an “unconventional spike” that is triggered by a large dendritic calcium transient has been proposed to be important(Terada et al., 2016). Indeed, a recent study also provided evidence that SK channels could regulate firing, because RNAi against SK was found to cause increases in the firing frequency of nociceptive neurons (Onodera et al., 2017). That study proposed that dendritically localized SK channels might respond to a dendritic Ca^++^ transient. Although our investigation of the localization of the SK channel proteins makes them well positioned to regulate firing of nociceptive neurons, our finding that SK is localized to axons is inconsistent with the hypothesis that SK is directly regulated by dendritic Ca^++^. Nevertheless, our comprehensive analysis, including the generation of null mutant alleles, and genomic and tissue specific rescue experiments, demonstrate a genuine involvement for SK channels in *Drosophila* nociception. Our studies of protein localization by a CRISPR engineered tagged SK channel, and anti-SK staining of an untagged rescuing cDNA suggest that the likely site of action for this important channel resides in the proximal axon segment and not in dendrites.

## Experimental procedures

### Fly strains

We identified K^+^ channel genes using the controlled vocabulary function at FlyBase (http://www.flybase.org) for all genes with K^+^ channel activity (http://www.flybase.org/cgi-bin/cvreport.html?id=GO:0005216). We used these fly strains: *w; md-Gal4;UAS-Dicer2*, *w; ppk-Gal4;UAS-Dicer2*, and *w; ppk-Gal4 UAS-GCaMP3.0*. Fly stocks with UAS lines and TRiP RNAi lines were provided by the Bloomington Stock Center. RNAi lines for screening were provided by the Vienna *Drosophila* RNAi Center (Dietzl et al., 2007). *Drosophila* stocks were raised on standard cornmeal molasses fly food medium at 25°C. Where possible, we used balancers containing the *Tb* marker so that the inheritance of the UAS insertion(s) could be followed. Exelixis lines, *SK*^*d01963*^ and *SK*^*f01403*^, were provided by the Exelixis collection at Harvard Medical School (Thibault et al., 2004) and used to generate *SK* mutant flies. *SK* genomic rescue strains were generated by injection of BAC CH321-90C05 (Venken et al., 2009) performed by Rainbow Transgenic Flies, Inc. for *PhiC31*-mediated chromosome integration into the *attP2* docking site. Transgenic flies containing the BAC rescue construct were crossed into *SK* deletion background to generate the genomic rescue strain.

### Molecular biology

To preliminarily screen for deletion of *SK*, polymerase chain reaction (PCR) was performed on genomic DNA extracted from wandering third-instar larvae with the putative FLP/FRT-mediated *SK* deletion. The Qiagen DNeasy^®^ Blood and Tissue kit for DNA extraction and the forward primer: 5’-ATGTCAATTCAGAAGCTTAACGACAC-3’, the reverse primer: 5’-TGCAGTTGCTGCTGATGCAGAT-3’ were used to show absence of *SK* gene product by PCR. Control primers targeting Caα1D, forward primer: 5’-CAACCGGATGTGAAGTGCG-3’ and the reverse primer: 5’-CTTGGCACTT CGCCTGAAGG-3’, were used to confirm integrity of DNA preps. Two additional primer pairs were used to confirm the SK deletion. From the upstream gene, *CanB*, into the pBacXP: forward primer 5’-GCAGTCGTGTATTTGCTGTcg-3’ and reverse primer 5’-TACTATTCCTTTCACTCGCACTTATTG-3’ (Primer pair 1). From the pBacWH into c-terminal end of *SK*, forward primer 5’-CCTCGATATACAGACCGATAAAAC-3’ and reverse primer 5’-ATCCTGGTGCCCTACGATCTATGT-3’ (Primer pair 4).

### CRISPR gene tagging

A V5 epitope tag was endogenously introduced at the SK locus (Figure S2A,B) using a single-stranded oligodeoxynucloetide (ssODN) homology directed repair CRISPR-Cas9 approach (Gratz et al., 2013). Briefly, the guide RNA sequence 5’-CCATTCAAGCGCCAACGCCC-3’ was cloned into the BbsI site of pU6-BbsI-chiRNA plasmid (Gratz et al., 2013) and co-injected at a concentration of 250ng/uL with the ssODN donor repair template 5’-GAGCGGATCGAGCAGCGGCGGAAC TTTTTACATCCTGACACAGCTGCAGTTGCCCCCATT<u>GGTAAGCCTATCCCTAACCCTCTCCTCGGTCTAGATTCTACG</u>CAAGCGCCAACGCCCCAATCGATGTTCAATGCAGCGCCCATGCTGTTTCCACATTCTAGG-3’ (IDT) at a concentration of 100ng/uL into *w*^1118^;; *PBac{vas-Cas9}*^*VK00027*^ embryos. Underlined sequence in the ssODN repair template indicates the newly introduced in-frame V5 epitope tag with an intrinsic XbaI restriction site (double underlined). Model System Injections performed embryo injections. F0 flies were crossed to *w*^1118^ and genomic DNA extractions were performed on single candidate flies for molecular screening of V5 integration events using PCR-RFLP analysis with the following primer pair: 5’-GAGCGTTTAACCAACCTAGAG-3’ and 5’-GCAGTTAGTGTTCGTCCAAAG-3’. PCR products showing the desired XbaI cleavage from single fly candidates were selected for further molecular verification of proper *SK::V5* integration (Figure S2B). All desired strains were appropriately balanced and fully sequenced at the genomic and transcript level (Figure S2C).

### Thermal nociception assay

Six females were crossed to three males. All crosses were kept at 25°C and 75% humidity on a 12 hour light cycle. Third-instar, wandering larvae were collected between days five to seven and stimulated with the noxious heat probe (6mm wide) controlled by the adjustment of voltage to temperatures of 42°C. We used a digital thermocouple (Physitemp, BAT-12) welded to the tip of the probe so that its temperature could be precisely measured, and delivered the stimulus by gentle touching of the larvae laterally, between or on abdominal segments four, five, and six. Each larva was only tested once. The experiments were videotaped; analysis of behavior was performed offline. The latency to response for each animal was measured as the time interval from the point at which the larva first comes into contact with probe until the larva initiates its first, complete (360° rotation) nocifensive response. For the initial screen, approximately 10-20 lines were tested per day and each day included a control cross for the driver line to the relevant isogenic background (*md-Gal4/+;UAS-Dicer 2/+*). We assigned alternative numeric labels to the VDRC lines so that the identity of the genes was blind to the tester throughout the entire screening process. The gene candidates were decoded following completion of the screen. Five to fifteen larvae were tested in the initial screen and an average latency was calculated. UAS-IR lines were selected for retesting if they should a latency that was one standard deviation from the mean seen in the control strains (*w; md-Gal4/+;UAS-Dicer2/+*, *w; ppk-Gal4/+;UAS-Dicer2/+*). We re-tested approximately 20-40 larvae for each line kept after the initial screen. For *SK* genetic mutant testing, only female wandering, third-instar larvae were tested.

### Mechanical nociception assay

Crosses were prepared as with the thermal nociception assay. Wandering, third-instar larvae were collected between days five and seven, and stimulated with calibrated (~ 30-50 mN) von Frey fibers as previously described (Hwang et al., 2007; Zhong et al., 2010; Mauthner et al., 2014). Noxious mechanical stimuli were delivered by the rapid depression and release of the fiber so that the fiber would begin to bend on the dorsal surface of the larva. The stimulus was delivered between abdominal segments four, five, and six. The percent response following the stimulus was calculated for each genotype. For *SK* genetic mutant testing, only female wandering, third-instar larvae were tested.

### Gentle touch assay

The gentle touch behavioral assay was performed as previously described (Kernan et al., 1994; Tsubouchi et al., 2012), wandering third instar larvae were collected as described for nociception assays, were stimulated with an eyelash fixed on the top of a opposite end of small paintbrush. The larvae were gently touched on their anterior segments during normal locomotion. Each larva was stimulated four times with the eyelash and an average touch score generated for each genotype. For *SK* genetic mutant testing, only female wandering, third-instar larvae were tested.

### Calcium imaging

Calcium imaging experiments were performed as described previously (Tsubouchi et al., 2012) but with a 50 mN probe.

### Immunostaining

Larvae were fileted in cold PBS, and then fixed for 30 minutes with 4% paraformaldehyde. Larvae were then washed with PBS-T, and placed in to a blocking buffer with 2% BSA and 10% normal goat serum at 4°C overnight. The next day, the filets were incubated at room temperature with the primary antibody for four hours, washed with PBS-T, and then incubated overnight at 4°C with the secondary antibody. Larvae were washed with PBS-T and then mounted in VectaShield or ProLong Gold anti-fade mounting medium. The primary *SK* antibody was kindly provided by the Dolph lab (Abou Tayoun et al., 2011). The SK antibody was used at a concentration of 1:2000, the mouse anti-V5 antibody (Invitrogen) was used at a 1:200 concentration, the GFP antibody was used at a concentration of 1:250, and the 22C10 antibody was used at a concentration of 1:100. All secondary antibodies were used at a concentration of 1:600 or 1:1000.

Brains were dissected from wandering third instar larvae in cold 1XPBS, fixed in 4% PFA for 30min, and washed 3×30 min in 1XPBS + 0.2% triton X-100 (PBS-T). Tissue was blocked in 5% NGS for 1 hr and allowed to incubate overnight at 4°C with the primary antibody (concentration of 1:200 mouse anti-V5 and 1:500 rabbit anti-GFP). Brains were then washed 3×30 min in PBS-T, allowed to incubate in secondary antibody (1:1000 AF-546 anti-mouse and AF-488 goat anti-rabbit) overnight at 4°C, washed 3×30 min in PBS-T, mounted in ProLong Gold anti-fade medium, and allowed to cure overnight at RT in the dark before confocal imaging.

### Confocal imaging

For live GFP imaging, larvae were anesthetized and mounted as previously described in (Mauthner et al., 2014). Confocal Z-stacks of the dorsal cluster from the SK^MI1037^ MiMIC line were taken on a Zeiss LSM 5 LIVE. For immunostained tissue, anti-SK images were taken on a LSM 5 LIVE confocal microscope using a 40X objective and anti-V5 images were taken on an LSM880 63x objective.

## Acknowledgements

We thank the Vienna *Drosophila* RNAi Center, Transgenic RNAi Project, and the Exelixis Collection at Harvard Medical School (NIH/NIGMS R01-GM084947). Stocks obtained from the Bloomington Drosophila Stock Center (NIH P40OD018537) were used in this study. We also thank Sarah Sweeny and Katherine H. Fisher for technical assistance and members of the Tracey laboratory for helpful discussion and assistance. This work was supported by grants from the National Institutes of Health, 5R21DC010222, 5R01GM086458 and 5R01NS054899 to W.D.T. Kia Walcott and Jessica Robertson were supported by pre-doctoral fellowships from the National Science Foundation.

## Supplemental Information

**Table S1.**
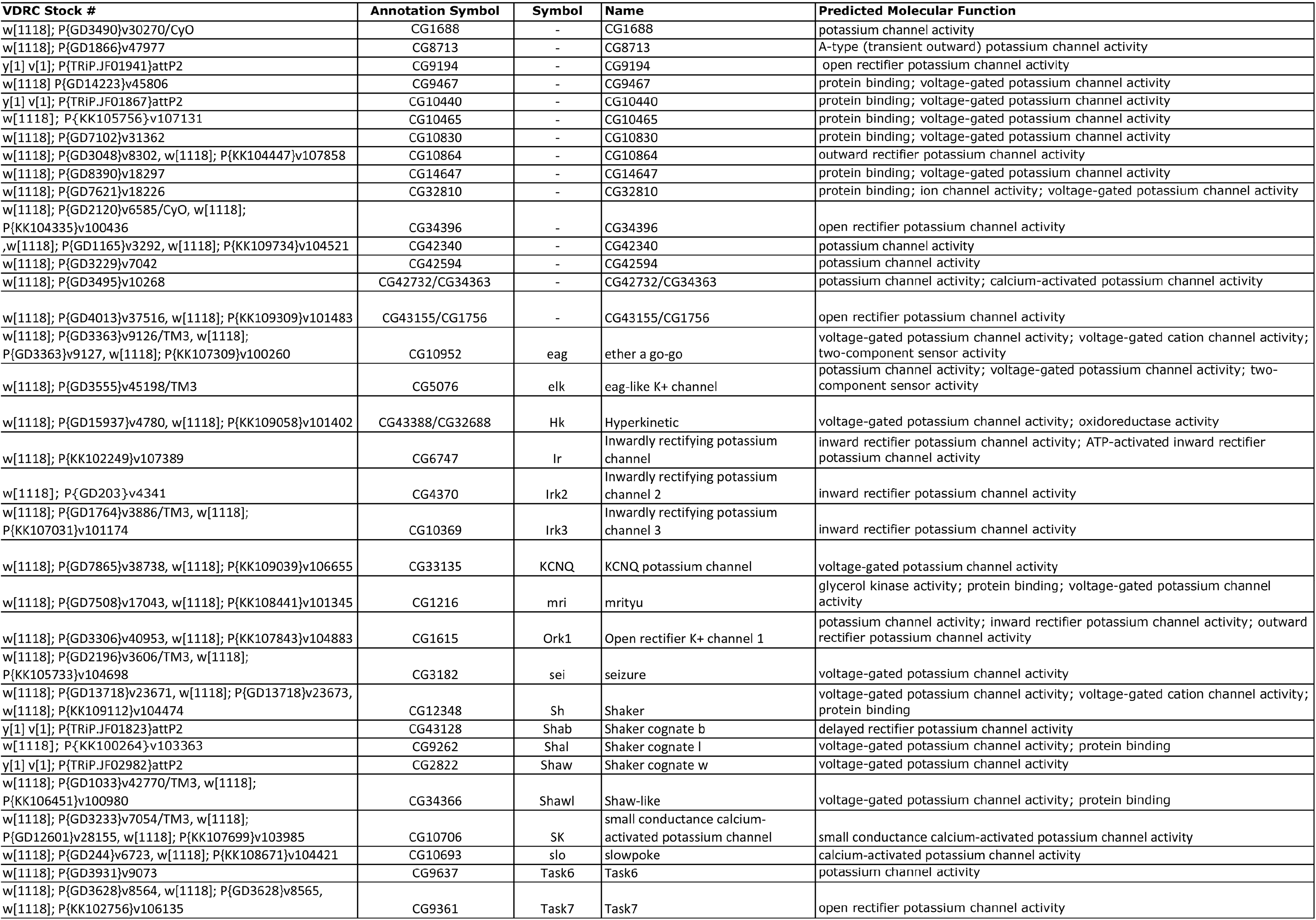
The complete list of K^+^ channel-targeting UAS-RNAi lines (n = 54) screened (with VDRC or TRiP stock #, annotation symbol, symbol, name, and predicted molecular function) is shown. In some cases, multiple RNAi lines were tested for a single K^+^ channel.

**Figure S1.**
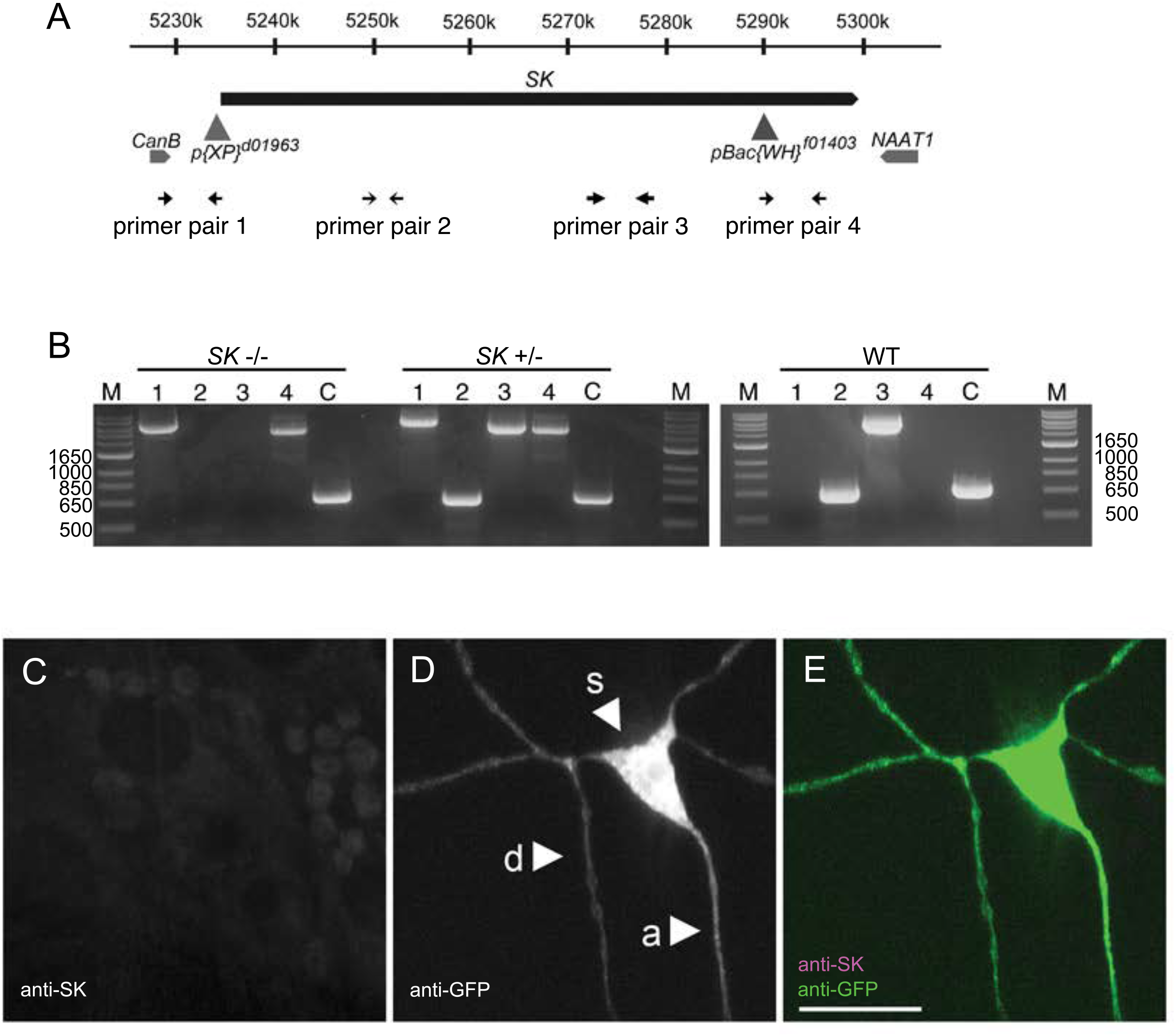
Confirmation of *SK* deletion by PCR. (A) The arrows numbered 1 - 4 represent distinct primer pairs for *SK* deletion confirmation by polymerase chain reaction (PCR). (B) Transgenic animals with *SK* deletion (*SK*^−/−^) show no product for the *SK*-specific primer pairs set 2 and 3. Transgenic animals heterozygous for the *SK* deletion (*SK*^+/−^) show products for all primer pairs as they have one copy of the WT and mutant chromosome each. WT transgenic animals (isogenic *white*) do not show products for primer set 1 and 4; they do not contain the transposable elements in their genome. Control primers against Caα1D demonstrate integrity of the genomic DNA preps. All transgenic animals used were virgin females (n>5). (D-F) Maximum intensity projection from z-stack images representative of the dorsal md neuron cluster (segment A4-A6) in larvae. Endogenous SK proteins are not at detectable levels in sensory neurons of *SK* null mutant larvae when immunostaining with anti-SK antibodies targeting the long isoform (genotype: *w SK^−^)*. SK protein expression (D and magenta in F) and class IV neuron (E and green in F).

**Figure S2.**
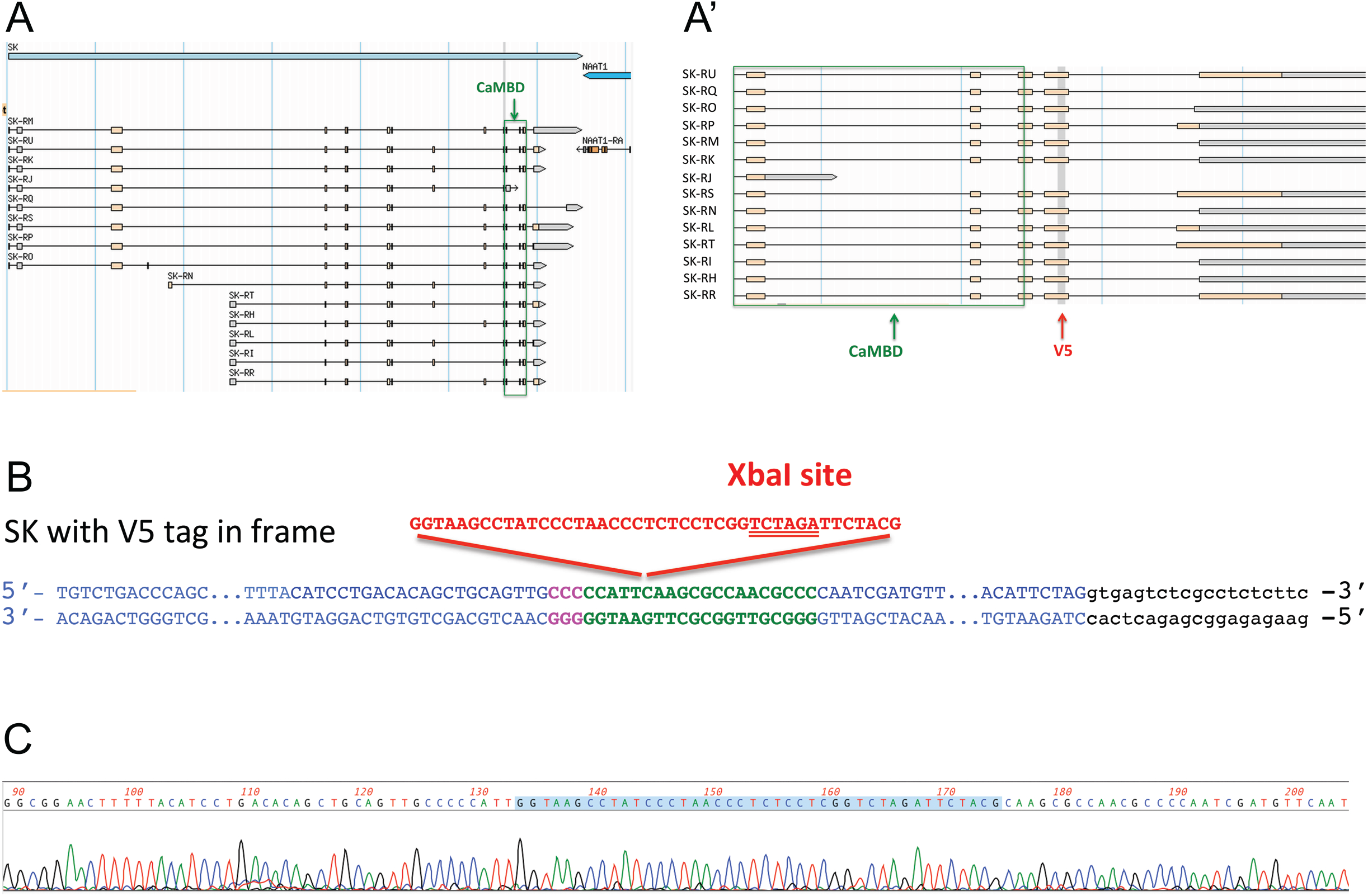
CRISPR engineered V5 epitope at *SK* locus. (A) Image of *SK* gene structure and predicted/known SK transcripts taken from Flybase version FB2017_04, released August 22, 2017 (Gramates et al., 2017). The exons encoding the calmodulin binding domain (CaMBD) in 13 out of the 14 predicted SK transcripts are highlighted in the green box. (A’) Zoomed in view indicating the in frame V5 epitope tag that was engineered into the SK locus using ssODN template (ssODN sequence in shaded grey box) for CRISPR mediated homologous repair. Predicted transcripts are noted and the V5 tag is inserted within last common exon after the CaMBD. (B) Predicted sequence of in frame V5 tag insertion site (indicated in red sequence) contains an XbaI restriction site (double underlined red sequence) for PCR-RFLP analysis. Exon sequence in black, intron sequence in blue, gRNA sequence in green, and PAM site in magenta. (C) Sanger sequencing results show proper insertion of V5 tag. Sequences are from cloned cDNA (generated by reverse transcription and amplified via PCR) targeting SK transcripts that span the V5 insertion and flank exon splice sites.

**Figure S3.**
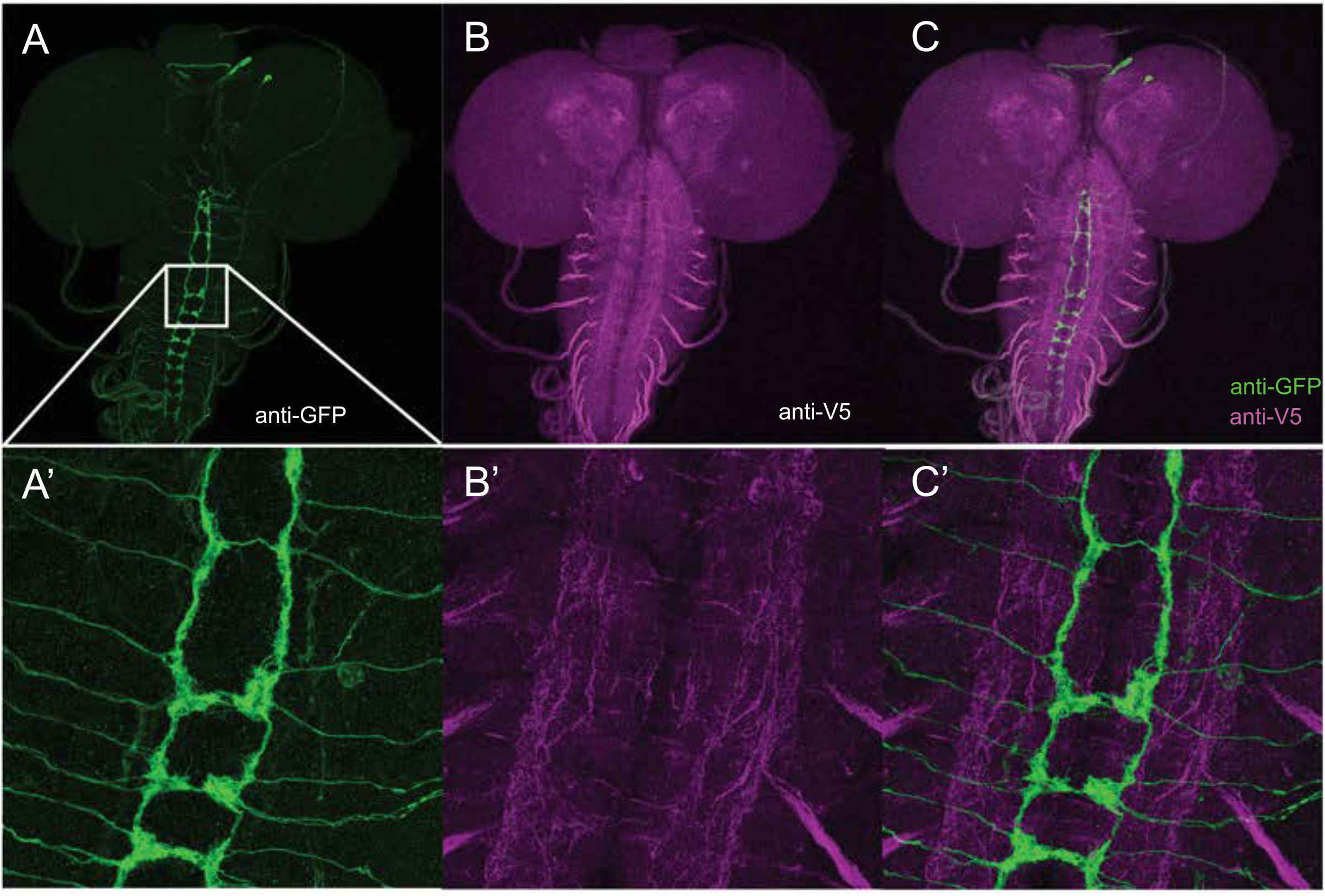
Immunostaining for SK proteins. (A-C). Maximum intensity projection from z-stack images representative of larval brains. SK::V5 proteins do not localize to class IV axon terminals in the ventral nerve cord (genotype: *SK::V5; ppk-GAL4 UAS-mCD8::GFP/+)*. Class IV projected terminals (A,C) and SK::V5 protein expression (B,C). (A’-C’) High magnification images of A, B, and C, respectively. d: dendrite, s: soma, and a: axon. Scale bars are 10μm.

